# Alleviation of salt stress in strawberries by hydrogen-rich water: physiological, transcriptomic and metabolomic responses

**DOI:** 10.1101/2024.07.25.605184

**Authors:** Renyuan Wang, Shaohua Chu, Dan Zhang, Xia Zhang, Yaowei Chi, Xianzhong Ma, Xunfeng Chen, Haiyan Yang, Wenjiang Ding, Ting Zhao, Yongfeng Ren, Xijia Yang, Pei Zhou

**Author notes:** Corresponding authors. Email addresses (Pei Zhou), (Xijia Yang).

## Abstract

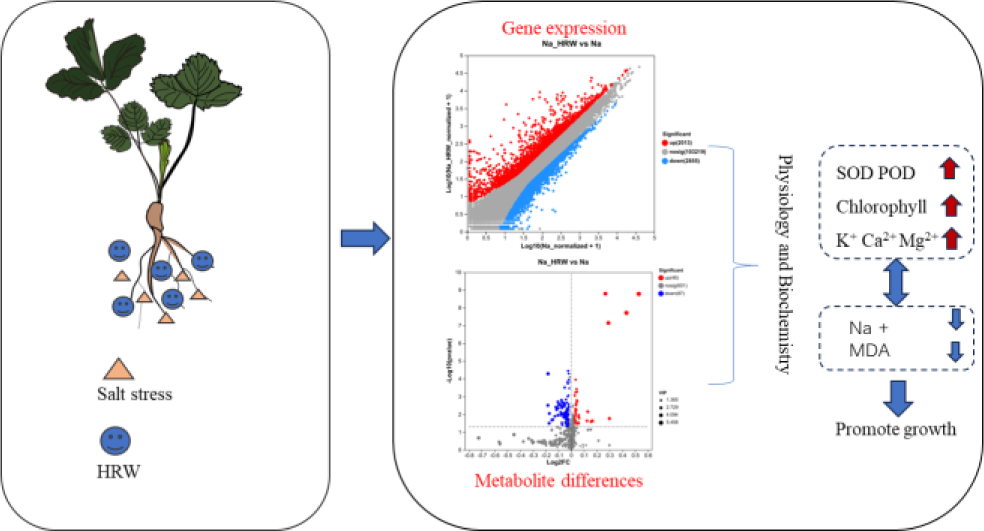

With the continuous advancement of climate change and human activities, cultivable land is becoming increasingly susceptible to the influence of salt ions. Therefore, it is crucial to mitigate the salt stress on plants through various approaches. In this study, strawberry seedlings were selected as the experimental plants, and a comprehensive transcriptomic and metabolomic analysis was conducted to elucidate the effects of hydrogen-rich water (HRW) on strawberries under salt stress. The results indicated that HRW significantly promoted plant growth, particularly increasing root biomass by 49.50%. Additionally, HRW regulated the levels of soluble sugars, malondialdehyde (MDA), and antioxidant enzymes, enhancing the cellular uptake of potassium ions and the expulsion of sodium ions. The levels of Ca^2+^ and Mg^2+^ in organelles increased by 2.06 and 2.45-fold, respectively. Transcriptomic analysis revealed that HRW substantially altered gene expression in strawberry roots; under salt stress, HRW up-regulated beneficial biological processes. Furthermore, genes related to ion absorption and transport, antioxidant enzymes, and cell wall biosynthesis were screened. Meanwhile, key common pathways were identified in differentially expressed metabolites (DEMs) and differentially expressed genes (DEGs) related to phenylpropane biosynthesis, alanine, aspartate, and glutamate metabolism, amino and nucleotide sugar metabolism, and galactose metabolism. A molecular mechanism for mitigating salt stress in strawberry seedlings by HRW was provided by the integrated approaches in this research, reflecting the potential applications of hydrogen gas in agriculture.

## 1. Introduction

Soil salinization is a significant environmental challenge worldwide, limiting plant growth, development, and crop yields. Climate change, industrial pollution, over-irrigation and improper use of fertilizers are the commonly existing main causes of soil salinization (Fu & Yang, 2023; Wang et al., 2023a). Meanwhile, approximately 1-2% of arable land is rendered unusable due to soil salinity issues each year (Bomle et al., 2021). When Na^+^ enters the cell, it will disrupt the intracellular water potential, causing osmotic stress. In addition, it also affects the balance of K/Na ions, leading to oxidative stress and ion stress (Das et al., 2023). However, roots are the pioneer organs that are exposed to salts in the cultivation (whether it is soil, substrate, hydroponics or fog cultivation) that have to adapt to the salinity of the cultivation substrate to sustain normal plant growth and promote water and nutrients absorption (Ji et al., 2013; Wang et al., 2021b; Zheng et al., 2024). Thus, it is necessary to study the salt adaptability of plants from the perspective of root systems. In addition, in order to cope with the growing population, agriculture has expanded to saline soils (Zulfiqar et al., 2022). It is crucial for people to explore strategies and methods to improve plant tolerance to salt stress in order to address food security challenges

Using exogenous substances is an effective method to improve plant tolerance to salt stress, including gas compounds like H_2_S, NO, as well as natural substances such as α-tocopherol, melatonin, trehalose, ascorbic acid, chitosan, and polyamines (Akram et al., 2022; Marone et al., 2022; Menhas et al., 2022). Compared to traditional exogenous substances, hydrogen gas has the greatest advantage of being green, non-toxic, and residue-free, which contributes to the promotion of green and sustainable development in agriculture (Li et al., 2021a). Currently, the primary method of delivering hydrogen in agriculture is through the use of HRW, which has been found that can enhance tolerance against different stresses in different plants (Wang et al., 2024; Zhao et al., 2021). The cultivated strawberry (*Fragaria × ananassa*, 8x) holds economic significance due to its nutritional richness and distinctive flavor profile (Mao et al., 2023; Song et al., 2024). However, strawberries are highly sensitive to salt stress (Mohammad et al., 2023). Thus, it is important to enhance the strawberry tolerance of salt stress.

Although many studies have been conducted on the application of hydrogen in plant stress, its specific mechanism is still unclear and requires further in-depth research and analysis. Therefore, revealing the intricate defense mechanisms of strawberries against salt stress at the metabolome and transcriptome levels in response to hydrogen is highly important. Transcriptomic technology has revolutionized the profiling of stress-responsive genes, while metabolomics represents a burgeoning scientific discipline born subsequent to genomics, transcriptomics, and proteomics (Gan et al., 2021; Song et al., 2022). It facilitates the systematic and quantitative analysis of an organism’s or cell’s global metabolome under external stimuli, acting as a pivotal link between the plant genome and its phenome. The synergistic analysis of metabolomics and transcriptomics promises enhanced precision in investigating biological phenomena, offering clearer understanding of the expression of key functional genes and the metabolic pathways they regulate (Lu et al., 2022; Mei et al., 2023; Wang et al., 2023b). Consequently, this approach will illuminate the molecular underpinnings and regulatory mechanisms governing plant stress resilience through the identification of key genes, metabolic pathways, and metabolites.

In this study, we hypothesized that under salt stress, hydrogen can activate specific pathways to alleviate salt stress on strawberries. To test this hypothesis, we utilize transcriptomic and metabolomic methodologies, focus on the root of strawberry seedlings, and thoroughly dissect the mechanisms through which HRW boosts salt tolerance in cultivated strawberries. This endeavor will significantly enhance our understanding of the molecular functionalities and regulatory mechanisms that underpin plant resilience to hydrogen-induced stress. Additionally, it will furnish theoretical and technical groundwork for integrating hydrogen into salt stress management in strawberry cultivation and beyond.

## 2. Materials and methods

### 2.1 Preparation of hydrogen-rich water

Bubbled pure hydrogen gas (99.99 %, v/v) generated by electrolysis from a hydrogen generator into 4000 mL deionized water at a rate of 320 mL min^-1^ for 3 h (QL-300, Saikesaisi Hydrogen Energy Co., Ltd., Shandong, China). Ensure hydrogen concentration over 1.0 ppm when HRW was used.

### 2.2 Plant materials and experimental design

Strawberry seedlings (*Fragaria×ananassa* Duch.‘Benihoppe’) used in the experiment were from tissue culture seedlings. After cultured the seedlings with substrate for one week, 5 leaves seedlings were chosen to a plastic pot (44*33*21cm) with 25 L soilless substrate. The experiment consisted of 2 treatments, 100 mM NaCl (NaCl) and 100 mM NaCl + 100% HRW (NaCl_HRW). Each treatments have 6 independent biological replicates. The experiment used a completely randomized block design, with 4 strawberry seedlings planted in each pot. To avoid the shock from salinity, the strawberry seedlings were subjected to a salinity treatment regimen in which they received 400 mL of NaCl solution at regular 48 h intervals, culminating in a total volume of 2800 mL over a span of 14 d. Concurrently, these samples were subjected to HRW treatment every two days. After the 21 d experimental period, collect plant samples and store them at −80 ℃.

### 2.3 Determination of biomass and chlorophyll content

Divided seedlings into shoot and root tissues, fresh weight was measured using an electronic balance. A chlorophyll assay kit was used to test the content of Chlorophyll (Nanjing Jiancheng Bioengineering Institute).

### 2.4 Determination of soluble sugar, MDA, and antioxidant enzyme activity

Crush the fresh samples that were previously stored under liquid nitrogen conditions (0.2g), and then 0.1 mol L^-1^ phosphate buffer solution was added, the mixture was ground to a homogenate, after centrifuged, and the supernatant was collected. MDA content was determined by thiobarbituric acid reaction method (Liu et al., 2024). Soluble sugar content and antioxidant enzyme activity were detected using a reagent kit (Nanjing Jiancheng Bioengineering Institute).

### 2.5 Determination of subcellular ion content

The subcellular components of strawberry seedling root cells were separated using a differential centrifugation method. Frozen root tissue (1g) was homogenized in a cold extraction buffer (250 mM sucrose, 50 mM Tris-HCl, 1 mM dithiothreitol) on ice. Subsequently, the homogenate was transferred to a centrifuge tube and centrifuged at 3500 g for 20 min. The precipitate was the cell wall component (F1); The supernatant was further centrifuged at 12000 g for 30 min, with the precipitate being the organelle component (F2) and the supernatant being the cell fluid (F3). All procedures were carried out at 4°C (Wang et al., 2021a). After extraction, the plant tissue was dried at 70°C. The dried tissue was digested at 120°C with a mixture of acids (HNO_3_ and HClO_4_, v/v, 3:1), diluted with ultrapure water, and analyzed using inductively coupled plasma emission spectrometry (ICP-OES, Optima 800, PerkinElmer, USA).

### 2.6 Transcriptomic analysis

Extract total RNA from plants using the OminiPlant RNA Kit (DnaseI) (Jiangsu Cowin Biotech Co., Ltd. China). Subsequently, the quality of RNA was assessed with 5300 Bioanalyzer (Agilent) and quantified by ND-2000. Construct a sequencing library using extracted high-quality RNA samples and perform sequencing analysis using Illumina Novaseq 6000 (Chen et al., 2022). Other methods and data analysis are described in the Supporting Information. Reference gene source: Fragaria x anassa; Reference genome version: WSU; Reference genome source: https://www.rosaceae.org/species/fragaria_x_ananassa/genome_v1.0.a1;

### 2.7 qRT-PCR validation analysis

18 candidate DEGs (9 upregulated and 9 downregulated) were screened for further qRT-PCR analysis. The detailed methods are shown in Supporting Information. Specific primers are listed in Table S3. Validation of the reliability of transcriptomic data is shown in Fig. S12.

### 2.8 Non-targeted metabolomic analysis

Weigh 50mg of fresh solid sample and place it in a 2mL centrifuge tube. Then, add 400 μ L of internal standard methanol extraction solution (methanol: water=4:1, v/v) containing 0.02 mg mL^-1^ L-2-chloroalanine and grind and extract with grinding beads. Grind the sample solution in a frozen tissue grinder at −10 ℃ and 50Hz for 6 minutes, and then extract it with low-temperature ultrasound at 5 ℃ and 40kHz for 30 min. Subsequently, the mixture was left at −20 ℃ for 30 min, centrifuged at 4 ℃ and 13000g for 15 min, and the supernatant was transferred to an injection bottle with an inner tube for analysis. Mix all sample metabolites of equal volume to prepare a quality control sample (QC) (Wang et al., 2022). Data analysis is shown in Supporting Information.

### 2.9 Statistical analysis

The data was analyzed using IBM SPSS Statistics 21.0 software (IBM, Armonk, USA). Statistical analysis was conducted using Student’s t-test, with a significance level set at *p* < 0.05.

## 3. Results and discussion

### 3.1 Effect of HRW on strawberry seedling growth response under salt stress

The effect of HRW on the growth of strawberry seedlings was clearly reflected through plant fresh weight and chlorophyll content. Photosynthesis was the physiological foundation for plant growth and development and was particularly sensitive to salinity stress. Hence, investigating how photosynthesis responded to salt stress was crucial for understanding the mechanisms by which plants cope with adverse conditions and enhancing their salt tolerance (Tobiasz-Salach et al., 2023; Weng et al., 2023). Chlorophyll was the primary pigment in photosynthesis, responsible for capturing light energy and converting it into chemical energy, which was fundamental to the process of photosynthesis. Therefore, the content of chlorophyll had a close relationship with the photosynthetic capacity of plants. Usually, a higher chlorophyll content indicated that plants possessed a stronger ability to capture and convert light energy, thereby potentially exhibiting higher photosynthetic efficiency (Cao et al., 2024; Helaoui et al., 2023). As shown in the Fig. 1a, HRW treatment significantly increased the fresh weight of strawberry seedlings, with an increase of 11.48% in the shoot and 49.50% in the root. Meanwhile, the application of HRW significantly increased the chlorophyll content of strawberry seedlings, with chlorophyll content increased by 25% and chlorophyll b content increased by 19.05% (Fig. 1b). This increased chlorophyll content could be seen as a manifestation of the enhancement of the strawberry seedling photosynthetic capacity and physiological adaptability in salt stress environments. MDA could be used to indicate the level of lipid peroxidation to some extent, while soluble sugars played a role in osmotic regulation (Chen et al., 2022; Lu et al., 2023b). As shown in Fig. 1 c, the MDA and soluble sugar content was significantly reduced. Furthermore, SOD and POD enzyme activity have significantly increased, and CAT and APX enzyme activity have no significant difference (Fig. 1 d and Fig S1). Those results illustrated that HRW could enhance specific enzyme activity to reduce oxidative stress under salt stress, and adjust content, thereby alleviating salt stress.

**Fig 1.**
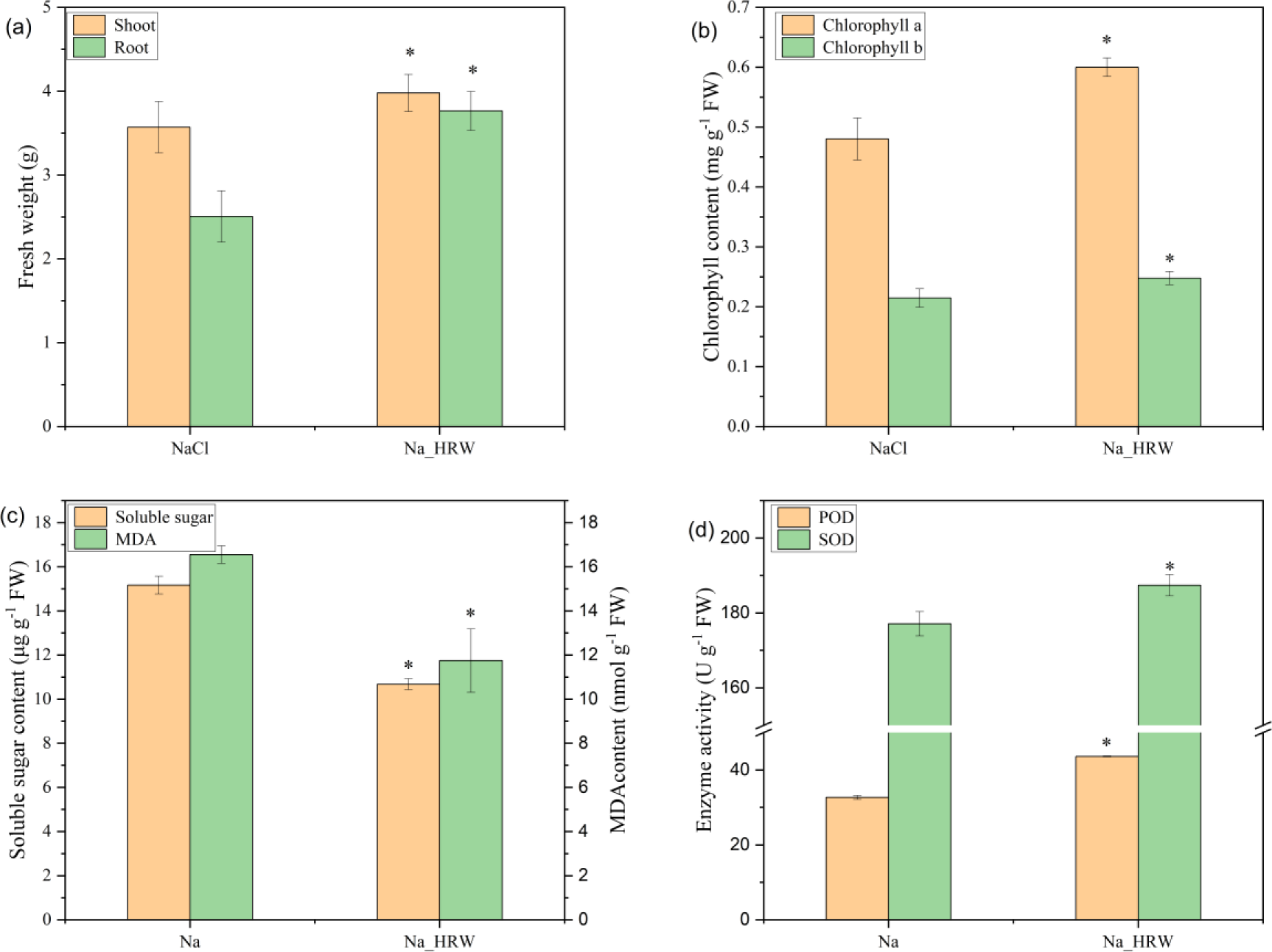
Effect of HRW on biomass (a), chlorophyll content (b), oxidative stress (c), and antioxidant enzyme activity (d) of strawberry seedlings under salt stress. The vertical bar indicates the standard deviation of three replications. For the same parameter, * above the bars denote a significant difference at *p* < 0.05. Na means 100 mM NaCl treatment group, Na_HRW means 100 mM NaCl with 1.0 ppm HRW treatment group.

### 3.2 Effect of HRW on ions subcellular distribution

As shown in Fig. 2, in the roots of strawberry seedlings, the content of K^+^ in F1 and F2 has significantly increased by HRW, increasing by 40.62% and 43.27% respectively, while in F3, K^+^ increased by 0.27mg g^-1^. Similarly, Ca^2+^ and Mg^2+^ were significantly increased in the F2 component, increasing by 2.06-fold and 2.45-fold respectively. Na^+^ was significantly reduced in all three components, by 7.93%, 16.79%, and 19.49%, respectively. The reason for the ion imbalance phenomenon by salt stress is that the ionic radii of K^+^ and Na^+^ are similar, and excessive Na^+^ competes with potassium channels in the apoplast. In addition, Na^+^ interferes with the function of K^+^, disrupting the stability of cell wall and membrane structure (Yamaguchi et al., 2013). Therefore, under salt stress, maintaining a potassium content within the cells above a certain threshold and keeping the K/Na ratio in the cells were key to the normal growth and salt tolerance of plants. In this study, Na^+^ significantly decreased in each component, while K^+^ significantly increased in each component, indicating that HRW reshaped the ion balance of inner cells and improved the K/Na ratio. These results are consistent with other studies (Wu et al., 2021)。 Reshaping the K/Na balance may restore root cellular function and integrity, which may be one of the reasons for alleviating salt stress.

**Fig 2.**
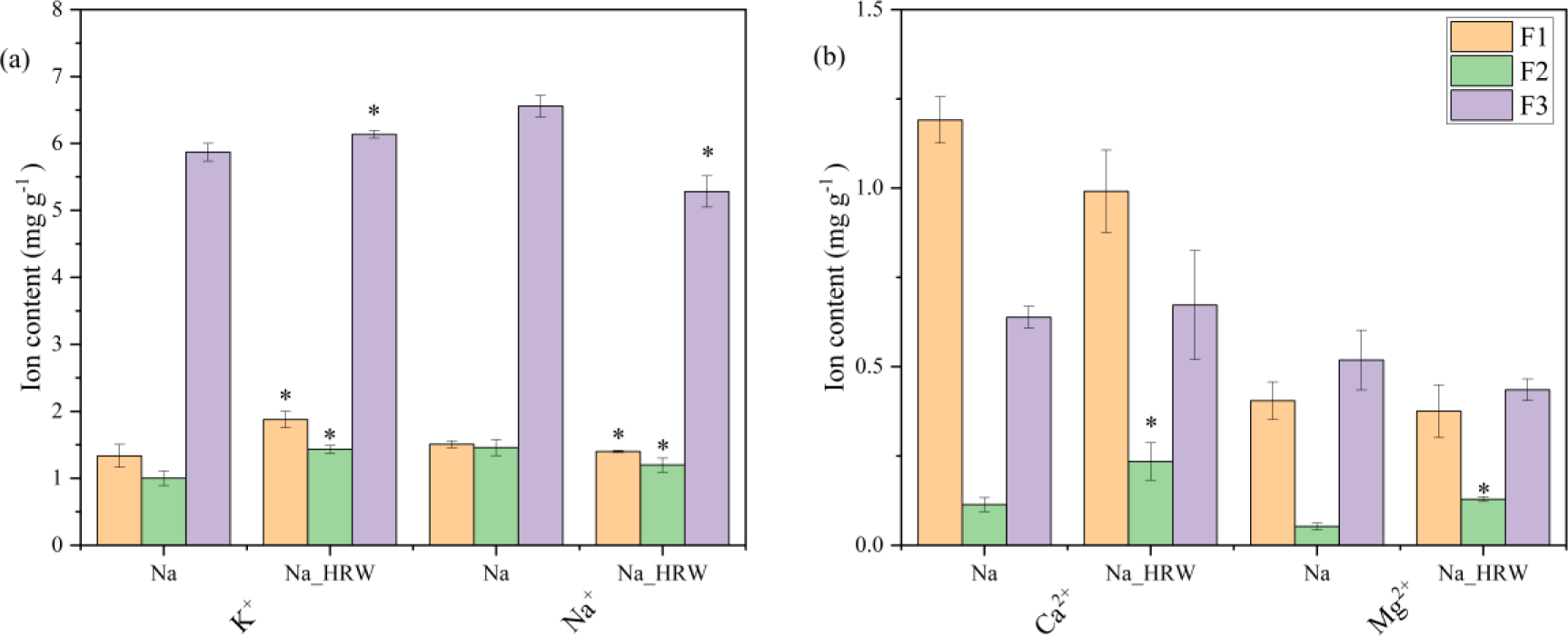
Subcellular distribution of ions. Na means 100 mM NaCl treatment group, Na_HRW means 100 mM NaCl with 1.0 ppm HRW treatment group. Subcellular components were divided into three components cell wall component (F1), organelle component (F2) and cell fluid (F3).

### 3.3 Transcriptome sequencing analysis

#### 3.3.1 Quality analysis of transcriptome sequencing

All samples exhibited Q20 and Q30 values exceeding 98% and 94%, respectively, confirming the reliability of the sequencing data in this study (Table S1). Uniquely mapped reads and total reads were above 75% (Table S2). The correlation heatmap displayed correlation coefficients above 0.99 for all treatments (Fig. S2), and PCA indicated significant expression differences among treatments (Fig. S3). Furthermore, |Fold Change| ≥ 2 and a corrected *p* < 0.05 were used as criteria for pairwise comparisons to identify differentially expressed genes (DEGs) between groups (Fig. 3a). Compared to NaCl treatment, HRW treatment identified a total of 4868 DEGs in strawberry roots, comprising 2013 up-regulated genes and 2855 down-regulated genes, suggesting that HRW modulated gene expression in strawberry roots.

**Fig. 3.**
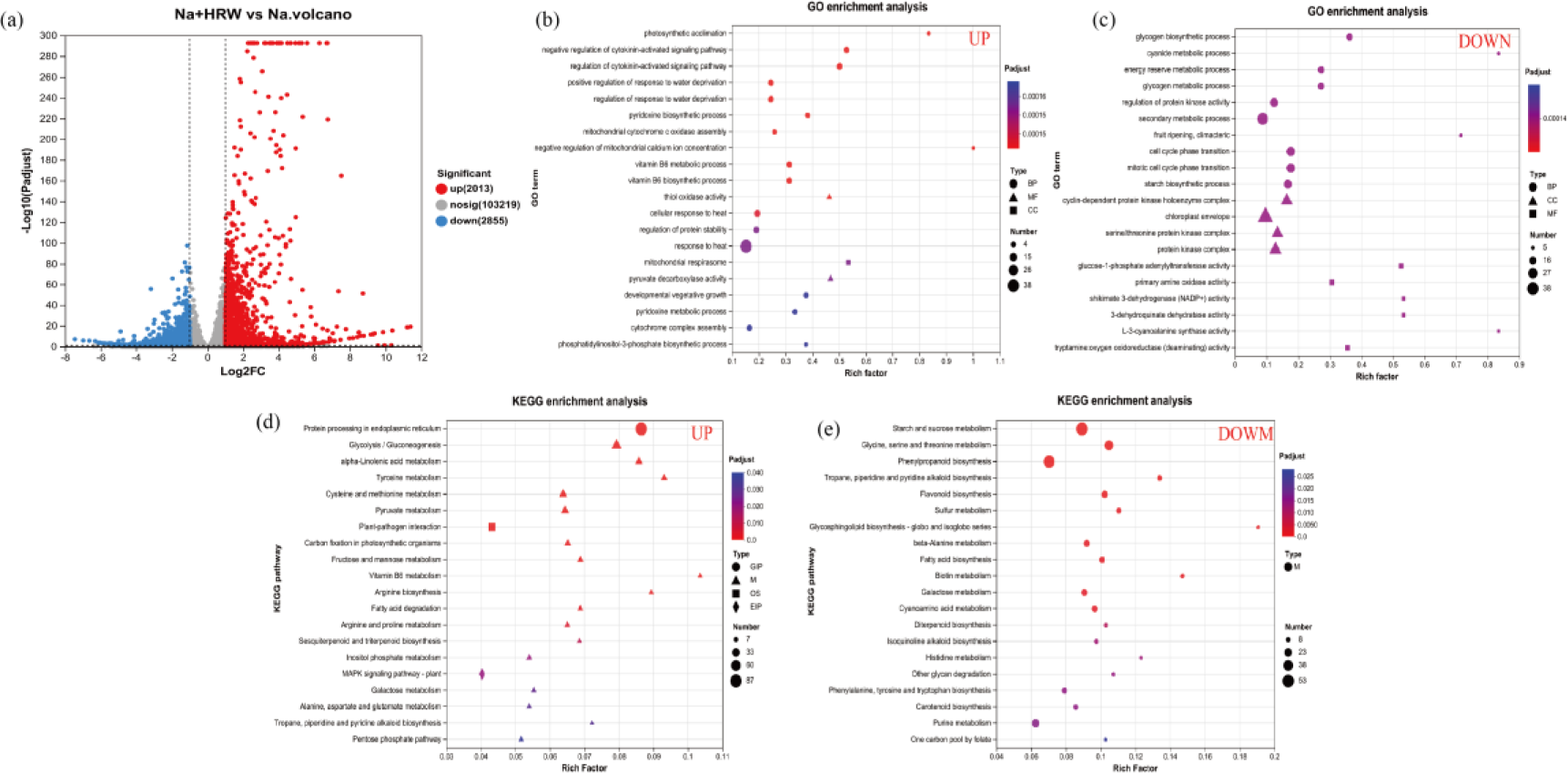
Transcriptomics analysis for different treatments. Volcano map of DEGs (a). Up-regulated DEGs enriched to GO analysis (b). Down-regulated enriched to GO analysis (c). Up-regulated DEGs enriched KEEG analysis (d). Down-regulated enriched to KEGG analysis (f). Na means 100 mM NaCl treatment group, Na_HRW means 100 mM NaCl with 1.0 ppm HRW treatment group.

#### 3.3.2 DEGs enrichment in GO analysis

GO analysis classified DEGs based on three ontologies: molecular function (MF), biological processes (BP), and cellular components (CC). The top 20 significantly enriched GO terms in the up-regulated DEGs shown in Fig. 3b included the most significantly enriched biological process (BP) term, “photosynthetic acclimation” (GO:0009643), “regulation of cytokinin-activated signaling pathway” (GO:0080037), and “regulation of response to water deprivation” (GO:0080036, GO:2000070). The molecular function (MF) term was “thiol oxidase activity” (GO:0016972), and the CC term was “mitochondrial respirasome” (GO:0005746). The impact of light adaptation on plants under salt stress included adjusting leaf anatomical structure, biochemical metabolism, chlorophyll and carotenoid content, mineral concentration, and overall plant biomass. These adjustments helped plants adapt to salt stress environments, and maintain optimal growth status, and photosynthetic efficiency (Zhang et al., 2020). The activation of photosynthetic adaptation might have increased the biomass and photosynthesis of strawberry seedlings. Similarly, cytoplasmic hormones such as cytokinin could be involved in plant growth and stress response (Zhao et al., 2023). As was well known, the accumulation of salt ions can lead to plant dehydration. The potential increase in plants water content and the compatibility of solutes could help maintain cell turgor pressure, create conditions for life activities, and slow down growth inhibition caused by water loss (Sanchez et al., 2012). In this study, the activation of regulation of response to water deprivation could have a positive effect on alleviating salt stress in strawberry seedlings. In the down-regulated DEGs (Fig. 3c), the BP terms were “glycogen biosynthetic process” (GO:0005978) and “cyanide metabolic process” (GO:0019499). The CC terms were “cyclin-dependent protein kinase holoenzyme complex” (GO:0000307) and “chloroplast envelope” (GO:0009941). The MF terms were “glucose-1-phosphate adenylyltransferase activity” (GO:0008878) and “primary amine oxidase activity” (GO:0008131). These results might have indicated that HRW regulated the reduction of energy consumption and maintenance of cellular functional balance in strawberry roots under salt stress, thereby reducing the generation of oxidative products that might have had harmful effects on cells. These could have been a pathway for HRW treatment to prevent excessive oxidation and reduce oxidative stress. In summary, these findings indicated that strawberry seedling gene expression responds sensitively to HRW. HRW activates genes in strawberries linked to critical biological functions such as photosynthetic acclimation, regulation of cytokinin-activated signaling pathways, and regulation of the response to water deprivation, etc. Thereby enhancing the tolerance of strawberry seedlings.

#### 3.3.3 DEGs enrichment in KEGG pathways analysis

To gain deeper insights into the effects of HRW on strawberry seedling biological processes, we also conducted an assignment of DEGs within the KEGG. As shown in Fig. 3d, the top 20 significantly enriched KEGG pathways in the up-regulated were “protein processing in endoplasmic reticulum” (map04141), metabolic processes (M) including “glycolysis / gluconeogenesis” (map00010), biological systems (OS) such as “plant-pathogen interaction” (map04626), and environmental information processing (EIP) like “MAPK signaling pathway - plant” (map04016).

The endoplasmic reticulum (ER) is vital for cellular homeostasis, participating in numerous processes such as folding and initial modifications of secretory and transmembrane proteins (Chen et al., 2020). Plants subjected to salt stress triggered ER stress, such as protein folding, protein modification, signal transduction, and ER-associated degradatio (Park & Park, 2019). A synergistic response by these ER proteins might have played a key role in plant defense against salt stress (Zhang et al., 2021b). Majority of these genes belong to the HSPs family. HSPs were believed to play an essential role in protein processing and protect plants from abiotic stress by preventing protein aggregation and misfolding. HSPs were closely associated with salinity tolerance (Cao et al., 2022; Wang et al., 2004). Additionally, the application of HRW activated the pathway of glycolysis/gluconeogenesis which played an important role. The salt-tolerant genotype sesame DEGs are related to glycolysis/gluconeogenesis pathways show a trend of enrichment (Zhang et al., 2019). Glycolysis/gluconeogenesis was significantly up-regulated by drought stress which could lead to the production of more adenosine triphosphate (ATP) derived from glycolysis in the root to adapt to adverse environmental conditions (An et al., 2016). Glycolysis/gluconeogenesis pathways were key pathways for energy production and carbon source supply. They decomposed and converted glucose to generate ATP and other energy molecules, and provided carbon skeletons for cellular metabolism. By upregulating the expression of relevant genes in these pathways, plants were able to increase the rate of energy production and carbon metabolism to meet the energy and carbon source demands of cells under salt stress. This regulation likely helped maintain cellular physiological balance and provided adaptability against salinity stress. In addition, the glycolysis/gluconeogenesis pathways were also associated with stress signal transduction and antioxidant reactions, potentially participating in the response and adaptation of plants to salinity stress through these pathways (Das et al., 2020; Zeng et al., 2019; Zhang et al., 2021a). Furthermore, the plant-pathogen interaction pathway also played a key role in alleviating the salt stress of HRW. Studies suggested that activating this pathway could enhance plant salt tolerance.(Fan et al., 2016; Peng et al., 2023; Zhang et al., 2021b). Moreover, the MAPKs play a crucial role in transmitting external signals such as mitogens, hormones, and various stresses in eukaryotes, and are beneficial for alleviating plant salt stress (Khan et al., 2023; Parihar et al., 2015; Xiong et al., 2020).

As shown in Fig. 3 d. The down-regulated DEGs, the top 20 significantly enriched KEGG pathways were all related to metabolic processes. The most significantly enriched pathways were “starch and sucrose metabolism” (map00500), “glycine, serine, and threonine metabolism” (map00260), and “phenylpropanoid biosynthesis” (map00260). HRW could have potentially enhanced energy for plant growth and development, antioxidant and protective mechanisms, secondary metabolite adjustment, and adaptation to salt stress by activating these pathways. In summary, these key DEGs revealed the potential molecular mechanisms by which HRW alleviated salt stress in strawberry seedlings.

#### 3.3.4 Screening of key DEGs

Bioinformatics analysis of DEGs revealed that HRW induced complex regulatory network pathway changes in root tissues. DEGs involved in ion regulation, antioxidant enzymes, and cell wall synthesis (Table S4). a total of 9 genes associated with sodium ion uptake and transport were screened, with 2 being up-regulated and 7 down-regulated. A total of 11 genes related to Na^+^ uptake and transport were identified, with 8 up-regulated and 3 down-regulated. Additionally, 13 genes related to Ca^2+^ were screened, with 9 up-regulated and 4 down-regulated. One gene associated with Mg^2+^ was up-regulated. In this study, 10 genes related to the antioxidant enzyme system were identified, including 5 genes related to POD (2 up-regulated, 3 down-regulated), 3 down-regulated SOD-related genes and 1 up-regulated CAT gene, and 1 down-regulated APX gene. A total of 41 genes associated with cell wall response to salt stress were identified, with 20 genes up-regulated and 21 down-regulated.

Balancing the K/Na ratio is an effective strategy for plants to cope with salt stress. However, high concentrations of Na^+^ limit the absorption of K^+^. HKT plays a crucial role in maintaining tomato K/Na homeostasis and salt tolerance (Li et al., 2021b). AKT1 and NHX2 plasma transporters play an important role in the absorption and transport of K^+^ by plants (Nieves-Cordones et al., 2014; Pinedo et al., 2015). In our study, 2 *NHX2* and *2AKT1* were identified among the up-regulated genes. Three *TPK1* and 2 *TPK5* were identified. Similarly, 2 Calcineurin B-like protein (*CBL*) and 3 Ca^2+^ uptake proteins were screened and all were up-regulated genes. Ca^2+^ signaling plays a key role in alleviating ionic stress by facilitating the extrusion or compartmentalization of Na^+.^ The absorption of Ca^2+^ also had a positive effect on plants in alleviating sodium ion stress (Mehra & Bennett, 2022), Therefore, we speculated that HRW regulated ion imbalance caused by Na^+^ accumulation in the roots of strawberry seedlings by regulating ion absorption and transport related genes.

Under stress conditions, plants were prone to accumulate excessive reactive oxygen species (ROS) in their tissues. Plants could effectively remove the excess ROS by activating the antioxidant enzyme system, thus avoiding oxidative damage to the plant (Chen et al., 2022). The research found that salt stress leads to a decrease in antioxidant enzyme activity, while the use of HRW increased the activity of SOD, POD, CAT, and APX (Wu et al., 2020). This finding was similar to our findings, indicating that HRW cloud enhance the activity of antioxidant enzymes to alleviate the salt stress experienced by strawberry seedlings.

Cell wall provided mechanical support and stability, enabling cells to resist external pressure and deformation. This was crucial for maintaining the integrity of cell morphology and structure in salt stress environments (Zhao et al., 2018). Meanwhile, cell wall underwent a series of changes under salinity stress, including adjustments in cell wall structure and composition. During salt stress, plants may increase the content of cell wall components like cellulose and pectin, potentially enhancing mechanical strength and tensile resistance of the cell wall, thereby contributing to the maintenance of cell integrity (Lu et al., 2023a; Rui & Dinneny, 2020). The plant cell wall is a highly responsive and dynamic structure that undergoes remodeling during plant growth and under biotic and abiotic stress, aiming to achieve adaptive regulation under various conditions. Some argue that the tolerance of plants to salt stress may be closely related to the formation of secondary cell walls and the deposition patterns of cellulose and lignin(Le Gall et al., 2015; Oliveira et al., 2020). In this study, the up-regulated genes mainly consisted of those encoding sucrose synthase, pectin methyl esterase, gluconate dehydrogenase, and expansin proteins. Plant cell wall expansion proteins played an important role in cell wall relaxation and expansin. They were widely recognized as key regulatory factors for cell wall extensibility response to stress and plant growth (Lu et al., 2013; Yang et al., 2023b). Thus, these results indicated that HRW could significantly regulate gene expression and metabolites related to the biosynthesis metabolism of strawberry seedling cell walls.

### 3.4 Metabolomics analysis

#### 3.4.1 Quality control of metabolomic data

This study utilized non-targeted metabolomics techniques (LC-MS) to explore various plant metabolites responsive to salt stress, with the aim of gaining a deeper understanding of how HRW affected the response mechanism of strawberry seedling roots under salt stress conditions. Quality control (QC) samples were well clustered together in the Fig. S4 that indicated the LC-MS equipment used in this study was stable. Meanwhile, under both positive and negative ion modes, the RSD < 30%, and peak accumulation exceeded 90%. These results indicate that the data is reliable. The OPLS-DA showed that Na group and Na_HRW group were clearly separated, indicating HRW changed metabolite profiles of strawberry root (Fig S5). In this study, a total of 601 metabolites were identified, among them which 135 differential metabolites (DEMs, VIP > 1 and *p* < 0.05) were screened (Fig. 4a). Compared with Na treatment, there were 48 up-regulated and 87 down-regulated metabolites in the HRW treatment. Among 135 DEMs, there were 33 Flavonoids (24.44%), 11 Carbohydrates (8.15%), 12 Glycerophospholipids (8.89%), and 6 Amino Acids (4.44%) (Fig S6).

**Fig. 4.**
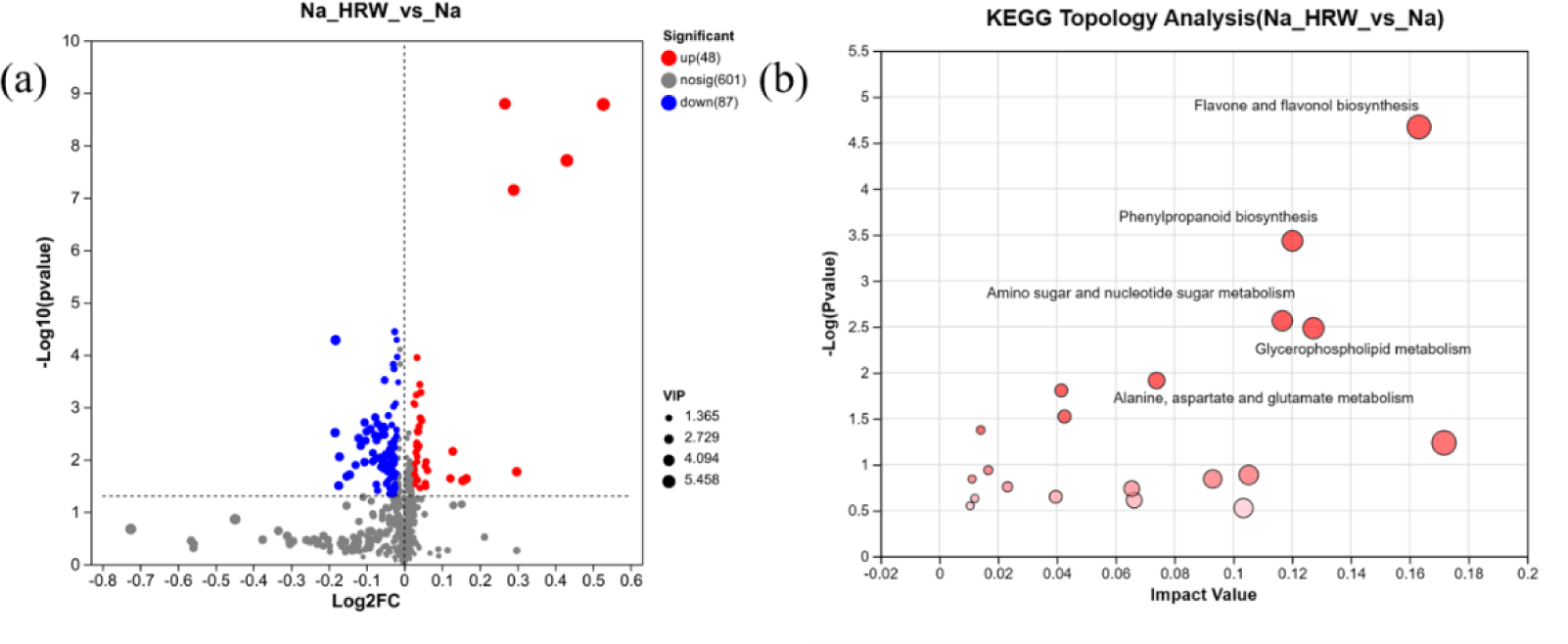
Volcano of DEMs (a). KEGG topological analysis (b). Na means 100 mM NaCl treatment group, Na_HRW means 100 mM NaCl with 1.0 ppm HRW treatment group.

#### 3.4.2 Key pathway analysis of DEMs

KEGG improved the query function for integrating metabolic pathways. Therefore, this study through topological analysis and metabolic pathway analyzed the enriched metabolic pathways in KEGG to investigate the key metabolic processes regulated by HRW in strawberry under salt stress. As shown in the fig. 4b, with the filtering criteria of *p* < 0.05 and the top 5 ranking based on the impact value, 5 significant enriched metabolic pathways were identified out of 36 metabolic pathways. These pathways were “flavonoid and flavonol biosynthesis”, “phenylpropanoid biosynthesis”, “amino sugar and nucleotide sugar metabolism”, “glycerophospholipid metabolism”, and “alanine, aspartate, and glutamate metabolism” (Fig 4b). Among them, 17 DEMs were enriched in 5 key metabolic pathways. The selected DEMs that were significantly up-regulated include phenolic acids and derivatives (ferulic acid, p-coumaryl alcohol, scopolin), flavonoids (astragalin, isoquercitrin), and amino acids (L-alanine).

#### 3.4.3 Analysis of DEMs in key pathways

Flavonoid and flavonol biosynthesis were considered one of the largest and most diverse groups of secondary metabolites in plants. It was believed that flavonoids acted as powerful antioxidants in plants, helping to neutralize free radicals and reduce oxidative stress damage to the plants. They were able to stabilize the redox balance within cells, regulate the synthesis and responses of auxins, and control important physiological processes, such as the growth of plant roots (Brunetti et al., 2013; Taylor & Grotewold, 2005). Similarly, under abiotic stress, flavonoid compounds were known to function as antioxidants by scavenging ROS, thereby preventing damage to cellular functionality and achieving an enhancement in plant tolerance (Izbianska et al., 2014; Jayaraman et al., 2021). In this study, compared with metabolites, the abundance of isoquercitrin and Astragalin was significantly increased after HRW treatment (Fig. 5b). Apigenin 7-O-glucoside and scolymoside abundance was significantly decreased (Fig S7a).

**Fig. 5.**
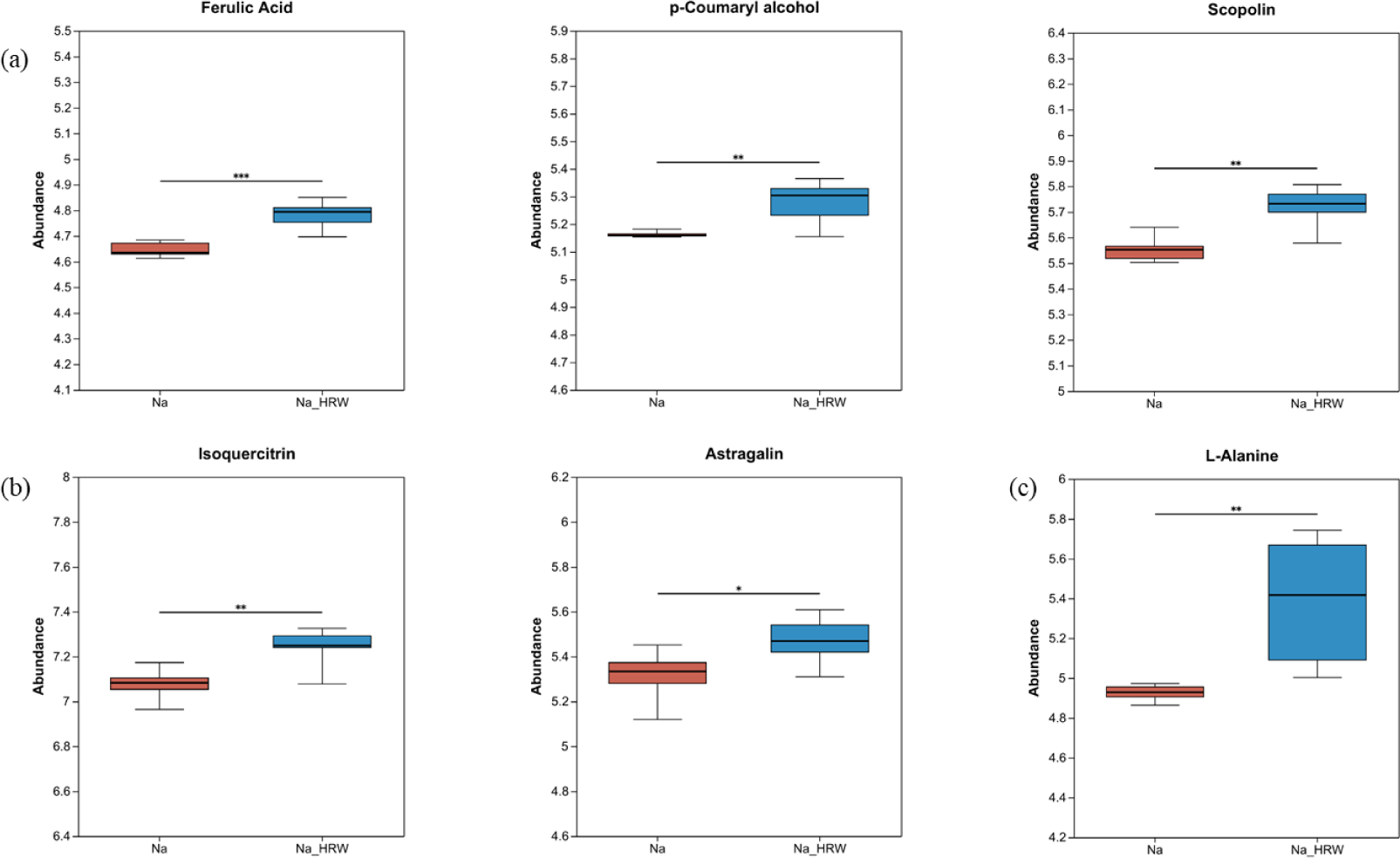
Significantly up-regulated phenolic acids and derivatives (a), flavonoids (b), and amino acids (c) in 6 identified DEMs. The * represents *p* < 0.05, ** represents *p* < 0.01, while *** represents *p* < 0.01. Na means 100 mM NaCl treatment group, Na_HRW means 100 mM NaCl with 1.0 ppm HRW treatment group.

Quercetin is a subclass of flavonoids involved in various plant physiological processes, such as seed germination, antioxidant mechanisms and photosynthesis. It was induced for the proper growth and development of plants, effectively providing plants with tolerance to a variety of biotic and abiotic stresses, which significantly enhances plant tolerance to various biotic and abiotic stresses by inducing normal growth and development (Singh et al., 2021). However, the research found that isoquercitrin showed higher activity levels than quercetin in superoxide anion scavenging assays, and isoquercitrin was more effective as a cell protective agent than quercetin (Li et al., 2016). Glycosylation is a significant modification process that plays a pivotal role in plant growth and various stresses. Astragalin, a glycosylated product of flavonols, was considered to potentially have a positive effect on improving plant resistance (Yang et al., 2023a).

As shown in Figure 5a, the abundance of ferulic acid, p-coumaryl alcohol, and scopolin significantly increased. Ferulic acid, a hydroxycinnamic acid derivative found in plant cell wall components, with diverse biological functions, it serves to neutralize excess ROS, directly scavenge free radicals, and inhibit enzymes generating free radicals. This compound has found extensive application in the investigation of various stresses (Zdunska et al., 2018). Research showed that exogenous addition of ferulic acid could enhance plant salt tolerance (Gupta & De, 2017). Different levels of salinity stress significantly induced the accumulation of FA in rice, concurrently reducing the accumulation of ROS and MDA. Similarly, pretreatment with exogenous FA could bolster cucumber resistance to dehydration stress by elevating proline and soluble protein content in the leaves, thereby reducing lipid peroxidation (Li et al., 2013). However, the significant decrease in the abundance of 5-Hydroxyferulic acid may be attributed to the significant antioxidant effects of HRW. These effects potentially promoted the activities of enzymes associated with FA synthesis and simultaneously inhibited the activities of enzymes responsible for converting Ferulic acid into 5-Hydroxyferulic acid, ultimately leading to a reduction in its abundance (Fig. S7b). P-coumaryl alcohol was a key intermediate product in the synthesis of lignin, which was ultimately converted into lignin by participating in multiple enzyme catalytic steps, affecting the synthesis and content of lignin. P-coumarin alcohol was involved in regulating the synthesis of lignin in pea sprouts and may have had an impact on the growth and stress resistance of pea sprouts (Lin et al., 2023). P-coumaryl alcohol was also considered an antioxidative compounds (Ly et al., 2003). Scopolin belonged to the coumarin class of compounds, which were coumarin derivatives related to clearing reactive oxygen species and defending against pathogens and osmotic stress could induce the accumulation of scopolin in roots (Beesley et al., 2023; Doell et al., 2018). Scopolin played a certain role in combating Cu toxicity and a low pH environment in lemon leaf slices (Zhang et al., 2022). Similarly, a decrease in scopolin could have affected the plant’s defense against pathogen infections, thereby weakening its resistance (Siwinska et al., 2014). The results showed that HRW regulated phenylpropanoid biosynthesis, increased certain antioxidant compounds, enhanced stress resistance, and alleviated the salt stress experienced.

The metabolic pathway of alanine, aspartate, and glutamate metabolism was able to regulate and protect the nitrogen metabolism balance, photosynthesis, and antioxidant reactions in plants. As shown in the figure, L-alanine was significantly increased (Fig. 5c) and L-asparagine significantly decreased (Fig. S7c). Studies showed that L-asparagine could not reduce the inhibitory effect of salt stress on the growth of rice seedlings (Lin & Kao, 1995), indicated that L-asparagine had a minor impact on plants under salinity stress. L-alanine was one of the smallest chiral compounds and was widely used in fields such as food, medicine, and veterinary science. However, L-alanine was able to promote the synthesis of chlorophyll, regulate the opening of stomata, and resist pathogens (Wu et al., 2023). In addition, L-alanine might have helped mediate the amino acid exchange between photorespiration amino acid metabolism and cytoplasmic amino acid metabolism (Betsche & Eising, 1986).

Abiotic stress activated the metabolic pathways of amino sugar and nucleotide sugar metabolism (Chen et al., 2023; Lu et al., 2023b). In this study, DEMs were also significantly enriched in these pathways. In addition, Numerous studies have confirmed that salt stress could induce plant phosphatidic acid (PA) accumulation. Similarly, salinity stress could induce phosphatidylcholine (PC) production (Li et al., 2024; Xue et al., 2024). Furthermore, PC stabilized the cell membrane by increasing the activity of phospholipase D and converted it into phosphatidylserine, phosphatidylethanolamine, and phosphatidylglycerol, thereby improving plant salt tolerance (Qiao et al., 2018). This study found that after the application of HRW, the content of PA and PC also significantly decreased, indicating that the salt stress was alleviated (Fig. S7e).

Overall, HRW altered root metabolism by producing more antioxidants and defense substances to maintain biological activity, thereby alleviating salt stress and promoting the growth of strawberry seedlings.

### 3.5 Joint analysis of transcriptome and metabolome

We conducted a combined analysis of transcriptomics and metabolomics to comprehensively elucidate the response of strawberry seedlings to salt stress alleviation under HRW, using the hypergeometric distribution algorithm to obtain pathways significantly enriched in gene sets and metabolites significantly enriched in metabolic sets. Four pathways were discovered through the study of two omics common pathways: phenylpropanoid biosynthesis, amino sugar and nucleotide sugar metabolism, alanine, aspartate and glutamate metabolism, galactose metabolism (Fig. 6a). There had a total of 5 metabolites and 64 genes involved in phenylpropane biosynthesis, while there were 4 metabolites and 42 genes in amino sugar and nucleoside sugar metabolism pathway. The pathway of alanine, aspartate, and glucose metabolism had 2 metabolites and 25 genes, while galactose metabolism had 2 metabolites and 29 genes.

**Fig. 6.**
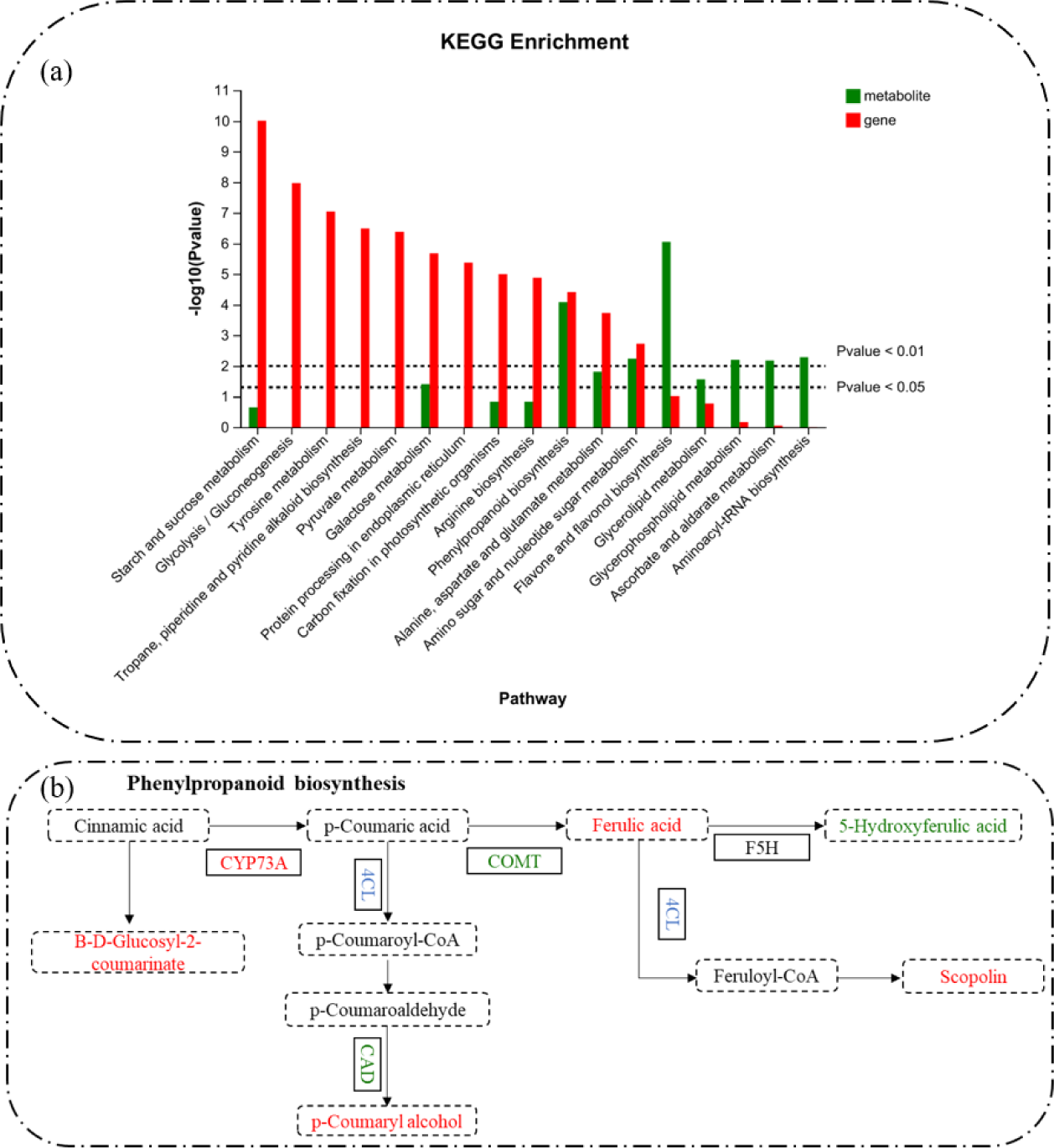
KEGG enrichment analysis of DEMs and DEGs (a). The main pathway of strawberry root system responds to salt stress under the action of HRW (b).

Phenylpropanoid biosynthesis is an important pathway which response to salt stress (Zhang et al., 2023). Products from the phenylpropanoid metabolic pathway contributed to the mitigation of ROS accumulation (Zhu et al., 2021). In this study, the metabolites in phenylpropanoid metabolic pathway that were elucidated also played a role in the clearance of ROS (Fig 5a). Similarly, phenylpropanoid metabolic pathway is a key pathway in the salt tolerance mechanisms of plants such as tomatoes, Sorghum, and barley (Ho et al., 2020; Jia et al., 2022; Ren et al., 2022). Galactose metabolism products may have been involved in the synthesis and repair of the cell wall, helping plants maintain the integrity of the cell wall and protect cell structures under salt stress conditions. The pathways of amino sugar and nucleotide sugar metabolism may have influenced energy metabolism pathways, regulating the energy supply and utilization of plants under salt stress, maintaining normal cell function. Salt stress greatly stimulated energy metabolism, while products from the pathway of amino sugar and nucleotide sugar metabolism was crucial for cell wall maintenance and repair (Lee et al., 2016). Transcripts and metabolites were co-enriched significantly in galactose metabolism and the amino sugar and nucleotide sugar metabolism pathway, indicating that HRW had activated those pathways. The decline in metabolite content in these pathways might have been due to HRW reshaping the ion balance, and alleviating osmotic stress, thereby affecting the products in the metabolic pathways.

## 4 Conclusion

In this study, our findings demonstrated that HRW effectively alleviated salt stress in strawberry seedlings. HRW not only promoted the growth of these seedlings under salt stress but also significantly increased chlorophyll content. HRW induced alterations in the levels of soluble sugars, MDA and antioxidant enzymes, thus improving the stress response capabilities of strawberry seedlings. Crucially, HRW reshaped the cellular K/Na ion balance and enhanced the absorption of Ca^2+^ and Mg^2+^ in the root cells. Through transcriptomic and metabolomic analyses, results indicated that HRW primarily affected the pathways of phenylpropanoids biosynthesis, alanine, aspartate, and glutamate metabolism, and carbohydrate metabolism. These pathways facilitated the production of more antioxidants and amino acids, which are essential for enhancing salt tolerance. Additionally, HRW might redistribute energy within the seedlings to further alleviate salt stress, thereby supporting their overall growth and survival under salt stress. Our research also involved screening genes related to ion regulation, antioxidant enzymes, and cell wall synthesis, revealing complex molecular interactions. Overall, our research provide fresh perspectives on the molecular pathways by which HRW mitigates the response of strawberry seedlings to salt stress and may offer theoretical support and technical assistance for using HRW in agricultural production.

## Author contributions

R.W. designed and participated in experiments and wrote the draft; S.C. and D.Z. designed the experiments; X.Z., Y.C. and X.M. conducted data analysis and image optimization; X.C., H.Y., W.D., T.Z. and Y.R. made critical revisions to the viewpoints presented in the manuscript; X.Y. and P.Z. designed the research and experiments and wrote and edited the manuscript.

